# SorLA restricts TNFα release from microglia to shape glioma-supportive brain microenvironment

**DOI:** 10.1101/2023.04.12.536447

**Authors:** Paulina Kaminska, Peter L. Ovesen, Mateusz Jakiel, Tomasz Obrebski, Vanessa Schmidt, Magdalena Bieniek, Jasper Anink, Bohdan Paterczyk, Michal Draminski, Anne Mette Gissel Jensen, Sylwia Piatek, Olav M. Andersen, Eleonora Aronica, Thomas E. Willnow, Bozena Kaminska, Michal J. Dabrowski, Anna R. Malik

**Affiliations:** Faculty of Biology, University of Warsaw, 02-096 Warsaw, Poland; Nencki Institute of Experimental Biology, 02-093 Warsaw, Poland; Max-Delbrueck Center for Molecular Medicine, 13125 Berlin, Germany; Institute of Computer Science, 01-248 Warsaw, Poland; Department of (Neuro)Pathology, Academic Medical Center, University of Amsterdam, 1105AZ Amsterdam, the Netherlands; Department of Biomedicine, Aarhus University, 8000 Aarhus, Denmark; Stichting Epilepsie Instellingen Nederland, 2103 SW Heemstede, The Netherlands

**Keywords:** VPS10P domain receptors, glioblastoma, intracellular sorting, phenotypic polarization, brain tumors

## Abstract

SorLA, encoded by the gene *SORL1*, is an intracellular sorting receptor of the VPS10P domain receptor gene family. Although SorLA is best recognized for its ability to shuttle target proteins between intracellular compartments in neurons, recent data suggest that also its microglial expression can be of high relevance for the pathogenesis of brain diseases, including glioblastoma (GBM). Here we interrogated the impact of SorLA on the functional properties of glioma-associated microglia and macrophages (GAMs). In the GBM microenvironment, GAMs are re-programmed and in turn lose the ability to elicit anti-tumor responses. Instead, they acquire glioma-supporting phenotype, which is a key mechanism promoting glioma progression. Our analysis of scRNA-seq data from GBM patients revealed that the pro-tumorigenic and pro-inflammatory properties of GAMs are linked to high and low *SORL1* expression, respectively. Using cell models, we confirm that SorLA levels are differentially regulated by the presence of glioma cells and by inflammatory cues. We further show that SorLA acts as a sorting receptor for the pro-inflammatory cytokine TNFα to restrain its secretion from microglia. As a consequence, loss of SorLA enhanced the pro-inflammatory potential of microglia, having a remarkable impact on glioma progression. In a murine model of glioma, SorLA-deficient mice develop smaller tumors and show hallmarks of anti-tumor response including altered microglia morphology, enhanced necroptosis, and massive neutrophil influx into the tumor parenchyma. Our findings indicate that SorLA is a key player in shaping the phenotype of GAMs, and its depletion can unlock an anti-tumor response.

**Significance statement:** Our study provides insight into the mechanisms shaping the tumor microenvironment in glioblastoma (GBM), the most prevalent and aggressive brain malignancy in adults. Poor prognosis in GBM largely results from the properties of the glioma milieu that blocks the anti-tumor response. We show that SorLA restricts release of the pro-inflammatory cytokine TNFα from microglia, thereby hampering their anti-glioma response. SorLA depletion reinforces the pro-inflammatory properties of tumor microenvironment and inhibits glioma growth. These findings have significant implications for our understanding of glioma biology, indicating SorLA-TNFα interaction as a potential target in GBM therapies. They also offer a new perspective on SorLA activities in microglia which emerge as highly relevant not only for the pathogenesis GBM, but also of other brain diseases such as Alzheimer’s disease.

## Introduction

Intracellular protein sorting is essential for maintaining cellular homeostasis and activity (1). Efficient sorting of proteins in the endocytic and exocytic routes not only ensures adequate levels of receptors and transporters on the cell surface, but it is also crucial for protein secretion (2). One of the key mechanisms governing intracellular sorting engages the VPS10P domain receptors. This receptors family entails five members (sortilin, SorCS1-3 and SorLA) which are mainly recognizedx for their roles in neurons (3). However, in a specific pathological context they can also be expressed in non-neuronal cells of the brain to shape their functional properties. For example, astrocytes activated after ischemic stroke express SorCS2, which is necessary for proper secretion of endostatin and for post-stroke angiogenesis (4).

SorLA, in humans encoded by *SORL1* gene, has been implicated in brain pathophysiology on genetic and functional levels. In particular, the protective role of SorLA in Alzheimer’s disease (AD) has been a subject of multiple studies, focusing primarily on its neuronal functions (3). However, recent data point to the potential context-dependent *SORL1* expression regulation in microglia. Thus, SNPs in *SORL1* previously associated with risk of sporadic AD possibly influence receptor gene expression in microglia rather than neurons (5). Microglial expression of *SORL1* was also noted in glioma patients’ brains (6). Still, potential microglial activities of SorLA in the diseased brain are unclear.

Microglia are innate immune cells of the brain that become activated under pathological conditions. Of note, in response to particular microenvironmental cues, microglia can enter various modes of activation and acquire diverse functional properties, ranging from pro-inflammatory to immunosuppressive (7). Although protecting the brain from insults seems to be the purpose of such activation, microglia can paradoxically also support disease progression. This is the case in glioblastoma (GBM), the common and most malignant primary brain tumor in adults (8). In GBM, massive accumulation of resident microglia, as well as of peripheral macrophages, is observed in the tumor microenvironment. These cells, collectively called glioma-associated microglia/macrophages (GAMs), share many functional properties including their tumor-supporting phenotype. GAMs account for up to 30% of tumor mass in human GBMs and in experimental gliomas (9–11) and their role in glioma progression and immunosuppression has been shown in various glioma models (12–14). Although they constitute a promising target in GBM therapy, the mechanisms shaping functional properties of GAMs are not fully understood.

Here, we hypothesized that SorLA may play a role in regulating the activities of GAMs by governing the intracellular sorting and secretion of its target proteins. Our *in silico* analysis indeed showed that *SORL1* expression in human GAMs is linked to their transcription profiles, likely reflecting diverse functional properties. Using cell models we further demonstrate differential regulation of SorLA expression by pro-inflammatory cues and by the glioma cells. Finally, we show that SorLA acts as a sorting receptor for TNFα to limit its release from microglia. Along these lines, tumor microenvironment in SorLA-deficient mice show enhanced pro-inflammatory properties which likely contribute to limiting glioma growth seen in these mice.

## Results

### SorLA expression levels in glioma-associated microglia/macrophages are linked to their activation mode

Activity of SorLA has been especially well documented in neurons, yet recent data suggest that it might also be expressed in other brain cell types in a specific pathological context. In particular, Abdelfattah et al. reported high expression levels of the *SORL1* gene in GAMs in human samples (6). In line with this notion, we could indeed detect SorLA in Iba1+ cells in human glioma sections, although not in all studied patient samples (Fig. 1A and Table S1). Induction of *Sorl1* expression in GAMs was recapitulated in a murine model of glioma (15). *Sorl1* expression in GAMs (CD11b+ cells) isolated from brain tumors 20 days after GL261 glioma cells implantation increased more than 3 times compared to the control cells. At the same time, SorCS2, another VPS10P receptor detected in this dataset, did not show such an induction (Table S2). Taken together, these data point to an intriguing hypothesis that SorLA levels in microglia/macrophages might be upregulated during activation of these cells towards a tumor-supporting phenotype.

**Figure 1.**
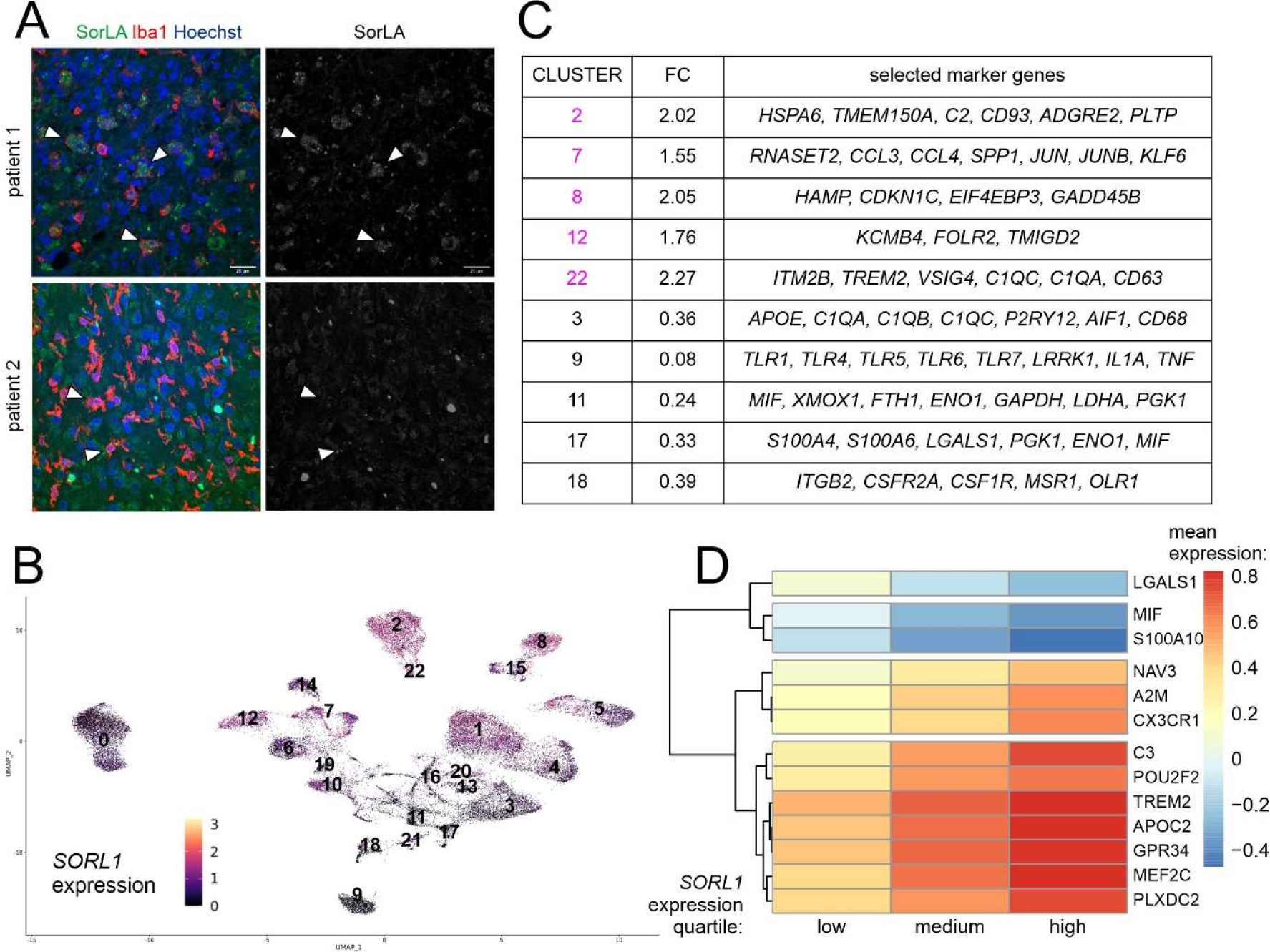
*SORL1* expression in human GAMs is linked to their functional properties. **(A)** SorLA is present in Iba1+ cells in some patients (#1) but absent from these cells in other cases (#2). **(B)** UMAP plot showing *SORL1* expression levels normalized with SCT, in clusters of human GAMs. **(C)** Selected marker genes of GAMs clusters with highest and lowest *SORL1* expression levels. FC – *SORL1* expression fold change; mean cluster expression against mean expression in remaining clusters. **(D)** Hierarchical clustering of the most significant genes based on their mean expression values. These genes were returned by MCFS-ID with the highest RI values and showed differential expression in the context of discretized values of *SORL1* gene expression.

To further link SorLA levels in GAMs to their functional properties, we analyzed scRNAseq data from human glioma samples published by Abdelfattah et al. (6), focusing on newly-diagnosed GBM samples (ndGBM). From this dataset, we selected 7 clusters that we classified as GAMs based on the expression of marker genes (*AIF1, CD68, ITGAM, P2RY12, TMEM119, CX3CR1*; Table S3 and Dataset S1). As reported by the authors of the study before, *SORL1* expression was limited to this GAMs population (Fig. S1). The only other cluster with noticeable expression of *SORL1* was characterized by expression of interferon-inducible genes (*IFI6, IFI27, IFITM1, IFITM2, IFITM3*) as well as endothelial markers (*EDN1, ABCG2, FLT1, PLVAP, PECAM1*) (Fig. S1 and Dataset S1). In all subsequent steps, we focused on the population of selected glioma-associated microglia/macrophages. These cells were pooled for further analysis to yield 22 GAMs clusters in which *SORL1* expression levels were evaluated (Fig. 1B). To shed light on the potential links between SorLA levels and the functional properties of GAMs we next investigated the marker genes of 5 clusters with highest and 5 clusters with lowest *SORL1* levels (Fig. 1C, Dataset S2). Among ‘high-SorLA’ clusters, cluster 7 was characterized by the expression of immediate early genes and *SPP1*, a gene associated with tumor-promoting GAMs (15), while cluster 22 showed high expression levels of several microglia/macrophages genes including *TREM2*, which was linked before to the pro-tumorigenic properties of tumor-associated macrophages in various cancers (16). At the same time, among ‘low-SorLA’ clusters, cluster 9 was characterized by expression of *TLR* genes as well as pro-inflammatory cytokines *IL1A* and *TNF*. Clusters 11 and 17 showed relatively high expression of a pro-inflammatory factor MIF and several glycolysis-related genes (*PGK1* and *ENO1* for both clusters, and additionally *GAPDH*, *LDHA* for cluster 11). Importantly, a metabolic switch towards glycolysis is a hallmark of pro-inflammatory activation of microglia/macrophages (17). Based on these results, we hypothesized that high and low SorLA levels might be associated with pro-tumorigenic and pro-inflammatory phenotypes of microglia/macrophages, respectively.

To gain deeper insights into the functional relevance of SorLA’s presence in GAMs, we performed Monte-Carlo Feature Selection (MCFS-ID) analysis (18, 19) on a single-cell level in the same dataset. MCFS-ID allows to determine features (here expression levels of a given gene) predicting the behavior of another feature (here, *SORL1* expression levels categorized to low, medium, or high). Among 25 top genes from the MCFS-ID differential levels of gene expression in the context of discretized values of *SORL1* were assessed (Fig. 1D and Dataset S3). This analysis highlighted microglia signature genes (*CX3CR1, A2M, C3*) and *TREM2* among the best predictors of high *SORL1* expression levels in GAMs. Moreover, high expression of a transcription factor MEF2C, known for its role in restraining microglial inflammatory response (20), was also linked to high *SORL1* transcript levels. Interestingly, we also noted a similar positive association between *SORL1* and *GPR34*, a microglial receptor required to arrest microglia in the homeostatic phenotype (21). Finally we identified *LGALS1, S100A10* and pro-inflammatory *MIF1* as predictors of low *SORL1* expression in GAMs.

Taken together, we propose a functional link between the activation status of microglia/macrophages and *SORL1* expression levels. In particular, our results point to the scenario where high *SORL1* expression occurs in tumor-supportive GAMs, while low *SORL1* expression is associated with pro-inflammatory phenotypes of microglia/macrophages.

### Loss of SorLA promotes TNFα release from cultured microglia

Since *SORL1* expression appeared related to the functional properties of GAMs, we tested whether SorLA levels might be specifically regulated by the cues triggering diverse microglial phenotypes. To study this phenomenon, we used primary murine microglia treated with LPS or co-cultured with glioma cell line GL261, an *in vitro* model to mimic pro-inflammatory stimulation and the impact of glioma-secreted factors, respectively. In line with our hypothesis, *Sorl1* expression increased in the presence of glioma cells, while it dramatically decreased upon LPS treatment (Fig. 2A).

**Figure 2.**
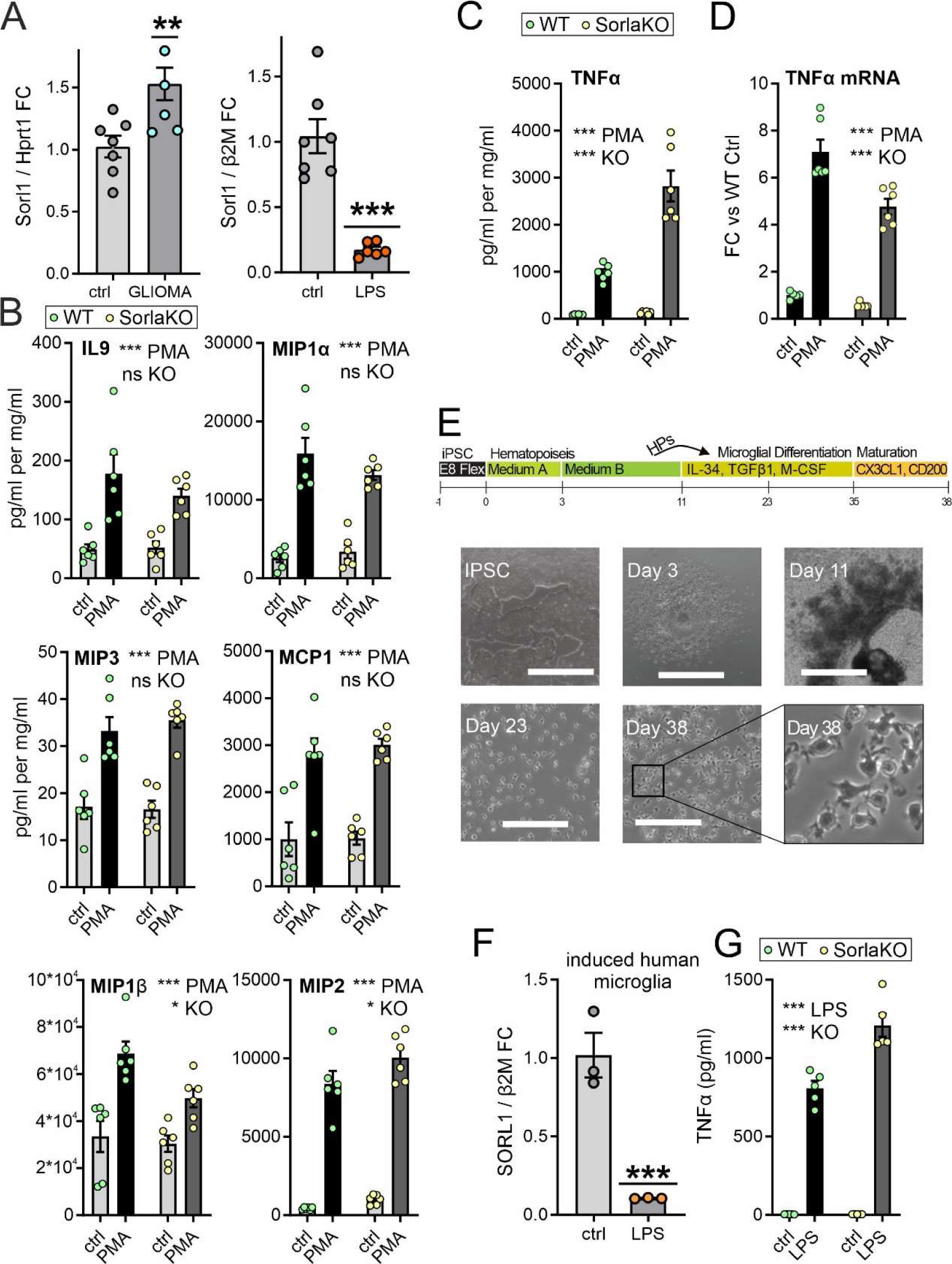
SorLA is differentially regulated by pro- and anti-inflammatory cues and restricts TNFα release from microglia. **(A)** *Sorl1* mRNA levels in primary murine microglia co-cultured with glioma cells (left) or stimulated with LPS (right) as assessed by qRT-PCR (relative to Hprt1 or β2M, respectively). Mean ± SEM is shown. n=5-7 independent microglial preparations. **, p < 0.01; ***, p < 0.001 in one-sample t-test compared to 1. **(B-C)** Cytokines levels as determined by ELISA in cell culture medium from primaryWT and SorLA-KO microglia either untreated (ctrl) or treated with PMA for 24 h. Cytokines levels were normalized to the protein content in the respective cell lysates. n=6 independent microglial preparations. Mean ± SEM is indicated. ns – not significant; *, p < 0.05; ***, p < 0.001 by two-way ANOVA. **(D)** TNFα mRNA levels in primary murine microglia stimulated with PMA as assessed by qRT-PCR (relative to β2M). Mean ± SEM. n=6 independent microglial preparations. *, p < 0.05; ***, p < 0.001 by two-way ANOVA. **(E)** Outline of the iPSC-to-microglia (iMG) differentiation protocol. Phase contrast images of iPSCs, HPs and iMG at different stages of the microglia differentiation. Scale bar, 1000 µm (iPSC, Day 3, day 11); 200 µm (day 23, day 38). **(F)** *SORL1* mRNA levels in human iMG stimulated with LPS as assessed by qRT-PCR (relative to β2M). Mean ± SEM. n=3 independent microglial preparations. **(G)** TNFα levels as determined by ELISA in cell culture medium from WT and SorLA-KO iMG either untreated (ctrl) or treated with LPS for 24h. n=5 independent iMG preparations ***, p <0 .001 by two-way ANOVA.

Distinct changes in microglial *Sorl1* expression seen upon activation by LPS and by the presence of glioma cells suggested that SorLA might be an active player in shaping functional properties of microglia. SorLA controls intracellular sorting of target proteins defining plasma membrane transport and secretion properties (22). As cytokines release from activated microglia is crucial for their activity and response to disease (7), we profiled cytokines released by WT and SorLA-deficient (SorLA-KO, SLKO) murine microglia upon PMA stimulation to uncover potential factors secreted in a SorLA-dependent manner. We did not observe any drastic alterations in cytokines secretion from SorLA-KO microglia as compared to control cells. Several cytokines were released in similar amounts in both genotypes, including IL-9, MCP1, MIP1α, and MIP3 (Fig. 2B). Secretion of MIP1β and MIP2 was slightly altered in SorLA-KO cells (decreased and increased, respectively). However, the most remarkable difference was seen in the secretion of the pro-inflammatory cytokine TNFα, which was released in higher amounts from SorLA-deficient microglia (Fig. 2C). This enhanced secretion of TNFα was not due to its increased expression, as mRNA levels were even downregulated in SorLA-KO cells (Fig. 2D). These results indicated that the alterations in TNFα release occur post-transcriptionally and might result directly from SorLA-dependent sorting mechanisms present in WTs, but absent in SorLA-KO microglia.

To further establish the relevance of our findings for human cells, we used microglia derived from induced pluripotent stem (iPS) cells either wildtype or genetically deficient for *SORL1* (Fig. 2E). We confirmed that LPS stimulation drives remarkable decrease in *SORL1* expression in human induced microglia (iMG; Fig. 2F). Moreover, loss of SorLA activity led to an increased TNFα release from iMG (Fig. 2G), indicating that SorLA-dependent control of TNFα secretion is conserved between the species and highlighting its potential relevance for disease pathogenesis.

### SorLA binds TNFα to control its intracellular trafficking

Typically, SorLA exerts its functions by binding target proteins and directing their intracellular trafficking. For example, this sorting activity of SorLA was documented for protein sorting between the TGN, endosomes and lysosomes, as well as in the recycling route via the Rab11+ compartment (23–25). To corroborate a potential role of SorLA in TNFα sorting, we first tested the colocalization of the two proteins in microglial cells. Indeed, immunostaining of PMA-stimulated BV2 cells revealed partial overlap of SorLA and TNFα signals (Fig. 3A). Next, we examined the interaction of SorLA with TNFα in co-immunoprecipitation (co-IP) assays. Towards this end, we overexpressed SorLA and GFP-tagged TNFα (or GFP alone) in HEK293 cells and pulled down the GFP tag. SorLA was present in the immunoprecipitate containing TNFα-GFP, while it was not visible in the control GFP-IP (Fig. 3B), supporting our notion that TNFα is a SorLA ligand.

**Figure 3.**
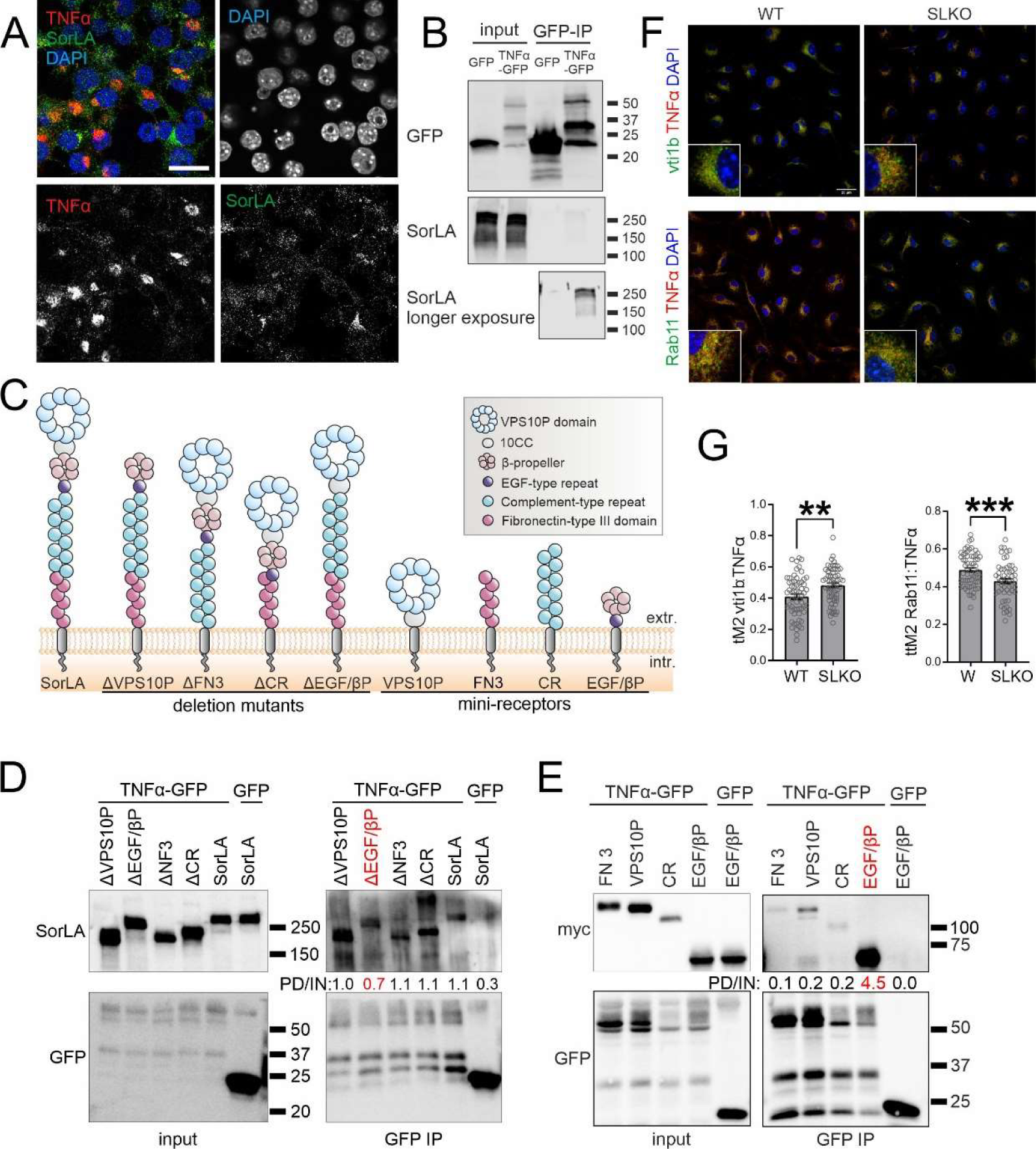
SorLA interacts with TNFα to regulate its subcellular distribution. **(A)** SorLA overlaps with TNFα in BV2 microglial cells stimulated with PMA. **(B)** Co-immunoprecipitation (co-IP) of SorLA with GFP-tagged TNFα overexpressed in HEK293 cells after GFP-IP. **(C)** SorLA variants used for co-IP experiments in D-E. **(D)** SorLA binding to TNFα is weaker for the ΔEGF/β-propeller mutant. **(E)** TNFα preferentially binds the EGF/β-propeller SorLA mini-receptor. **(F)** Colocalization of TNFα with markers of recycling endosome (Rab11) and secretory vesicles (vti1b) is altered in SorLA-KO microglia. **(G)** Thresholded Manders coefficients (tM) calculated for colocalization of Vti1b and Rab11 with TNFα.

As the structure of SorLA entails several domains capable of cargo binding (Fig. 3C), we sought to identify the domain responsible for the interaction with TNFα. Using deletion mutants lacking particular SorLA domains (Fig. 3C) in our co-IP experiments, we observed that a SorLA variant devoid of the EGF-type repeat and the β-propeller (ΔEGF/βP mutant) showed the weakest binding to TNFα as compared to the other mutants and the full-length receptor (Fig. 3D). In an alternative approach, we used myc-tagged mini-receptors composed exclusively of particular SorLA domains (Fig. 3C). In line with our prior observations, the most efficient co-IP with TNFα was noted for the mini-receptor encompassing the EGF-type repeat and β-propeller (EGF/βP, Fig. 3E). In summary, SorLA binds TNFα predominantly via an extracellular motif containing EGF-type repeat and β-propeller.

Finally, we elucidated the impact of SorLA on the intracellular trafficking of TNFα. We analyzed its colocalization with the markers of subcellular compartments in WT and SorLA-deficient primary microglia stimulated with PMA. We did not observe the TNFα presence in lysosomes nor late endosomes (stained with anti-Lamp1 and anti-Rab7 antibodies, respectively) in any of the genotypes (Fig. S2). Rather, TNFα was colocalizing with the Golgi (GM130-positive structures) and Rab11+ recycling endosomes, as well as with Vti1b, known for its important role in the TNFα secretory route (26). SorLA deficiency did not affect the presence of TNFα in the GM130+ compartment (Fig. S2), but it increased the colocalization of Vti1b with TNFα and caused a concurrent loss of TNFα from Rab11+ endosomes (Fig. 3F-G). These results indicated that loss of SorLA shifts the trafficking of TNFα towards the secretory pathway, which could explain the increased TNFα release from SorLA-KO microglia.

### Loss of SorLA limits glioma growth, promotes inflammation and necroptosis

SorLA emerged as a critical factor controlling the functional properties of microglia, restricting their pro-inflammatory activities. Moreover, in glioma patients, *SORL1* expression levels in GAMs were related to their transcription profiles. We further speculated that the presence of SorLA might have an impact on the functional properties of GAMs and, consequently, on tumor microenvironment and glioma progression.

Thus, we asked whether SorLA presence in the host cells has impact on glioma development in a murine model. Glioma GL261 cells carrying luciferase and tdTomato transgenes were implanted to the striata of WT and SorLA-KO mice and the tumor growth was followed for 21 days. Using this model, we showed that glioma growth is limited in SorLA-deficient mice (Fig. 4A-B), supporting our notion that the presence of receptor in host cells is critical for establishing a tumor-promoting microenvironment. As SorLA restricted TNFα release from primary microglia, we next verified whether it also has impact on the inflammatory responses in glioma-bearing brains *in vivo*. Since TNFα was undetectable in our samples, we evaluated downstream effects of the TNFα activities and the hallmarks of a pro-inflammatory phenotype of microglia.

**Figure 4.**
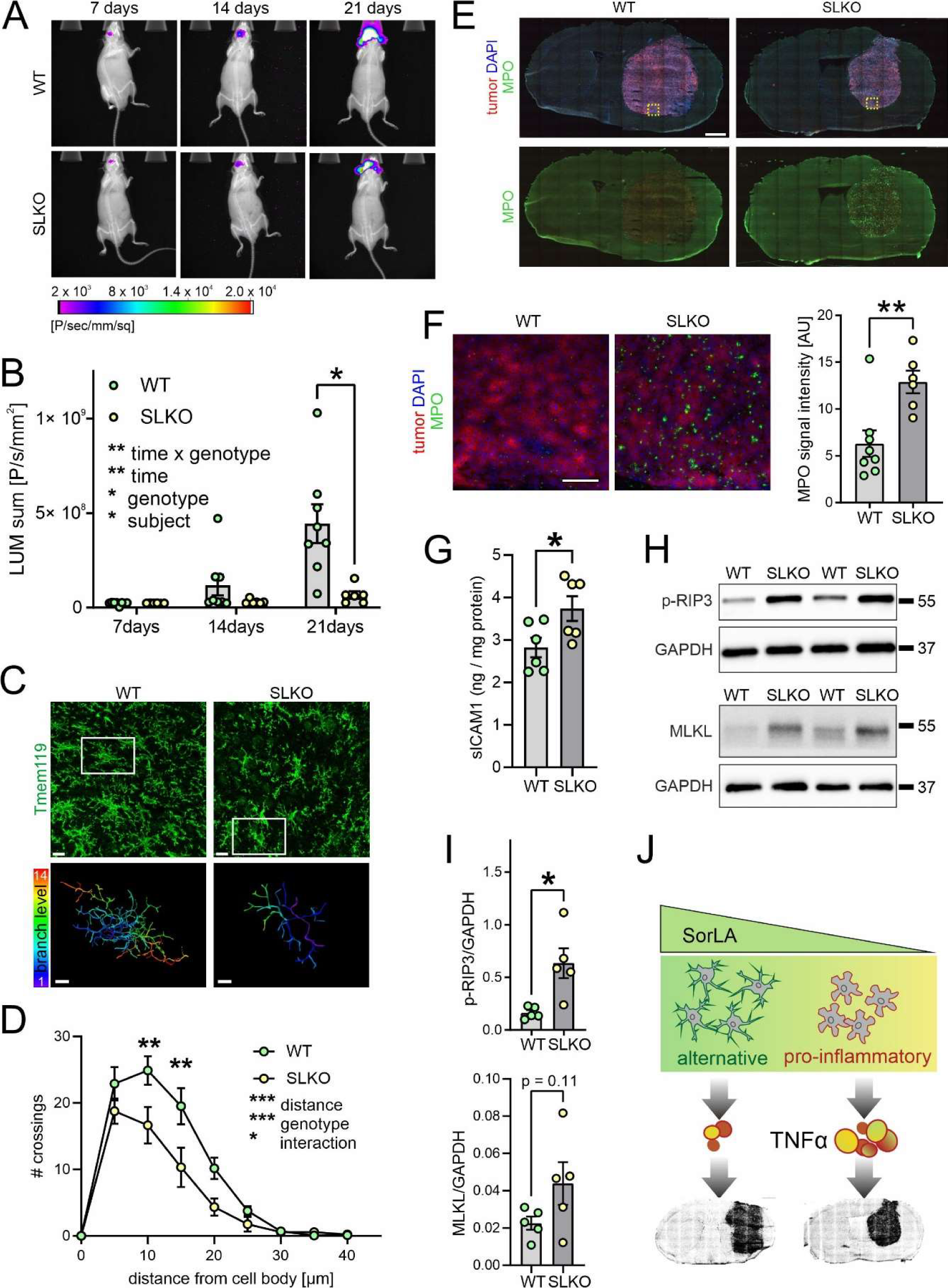
SorLA deficiency restores anti-tumor brain microenvironment. **(A)** Representative images of bioluminescence signals emitted by luciferase-expressing gliomas in WT and SorLA-KO mice at 7, 14 and 21 days post-implantation. Relative signal intensities represented by color are combined with X-ray images. **(B)** Bioluminescence signals measured as in (A) at indicated days post-implantation. Mean ± SEM is indicated. *, p<0.05 in repeated measures two-way ANOVA with Sidak’s multiple comparisons test; n=6-8 mice per genotype. **(C)** Upper panel: representative images of microglia morphology revealed by Tmem119 staining in WT and SorLA-KO in glioma margin 21 days post-implantation. Scale bar, 15 µm. White box indicates the cell reconstructed below. Lower panel shows reconstructed microglia branches; color depicts branch level. Scale bar, 5µm. **(D)** Sholl analysis of microglia morphology reconstructed as in (C). Mean ± SEM is indicated. ***, p<0.001; **, p<0.01; *, p<0.05 in two-way ANOVA with Sidak’s multiple comparison test; n=4 mice per genotype; for each mouse 5 cells were quantified and an average of obtained values was treated as an individual data point. **(E)** Representative images of murine brains sections 21 days post-implantation stained for neutrophils marker MPO and counterstained with DAPI (blue). Scale bar, 1 mm. Yellow box indicates the area imaged with higher magnification in (F). **(F)** MPO+ cells (green) in the glioma mass (red) in WT and SorLA-KO brains. Scale bar, 100 µm. Right panel: quantification of MPO signal intensity observed in WT and SLKO mice. Mean ± SEM is indicated. n=6-8 mice per genotype. **, p<0.01 in unpaired two-tailed t-test. **(G)** Levels of sICAM1 in soluble fractions extracted from glioma-bearing brain hemispheres 21 days post-implantation, normalized to protein content. Mean ± SEM is indicated. *, p<0.05 in unpaired two-tailed t-test; n= 6 mice per genotype. **(H)** Western blot analysis of necroptosis markers, p-RIP3 and MLKL in WT and SorLA-KO glioma-bearing hemispheres at 21 days post-implantation. Detection of GAPDH was used as a loading control. **(I)** Quantification of the western blot analysis as in (H). Signal intensities for MLKL and p-RIP3 were normalized to GAPDH signal. Mean ± SEM is indicated. *, p<0.05 in unpaired two-tailed t-test; n= 5 mice per genotype. **(J)** Schematic representation of the proposed mechanism. Low SorLA levels in GAMs promote release of TNFα and enhance anti-tumor responses.

The microglia activation status is reflected by the changes in their morphology (27–29). In essence, the tumor-supportive phenotype is characterized by ramified morphology with longer and more branched processes. By contrast, pro-inflammatory microglia present compact morphology and shorter extensions. These features can be evaluated by Sholl analysis, which quantifies the number of processes crossing the spheres of increasing radius, centered at the cell soma. This analysis performed on the tumor-surrounding Tmem119+ cells revealed remarkable differences between WT and SorLA-KO microglia (Fig. 4C-D). In detail, SorLA-KO microglia showed a compact morphology with shorter processes, which can be attributed to its more pro-inflammatory phenotype. By contrast, a branched morphology of microglia was seen in WTs that could correspond to a homeostatic or tumor-supporting state. These differences were also reflected by an apparent global decrease in Tmem119 signal in the tumor-surrounding tissue in SorLA-KO mice (Fig. S3). Of note, in the contralateral glioma-free hemisphere we did not observe any genotype-dependent differences in microglia morphology (Fig. S4).

Another important consequence of the inflammatory response is the influx of peripheral immune cells into the affected tissue. Locally released factors attract circulating leukocytes and promote their migration, eventually driving their infiltration into the inflamed area. In our glioma model, the infiltration of galectin-3+ macrophages and CD8+ T-lymphocytes into the tumor mass was similar for both WT and SorLA-KO mice (Fig. S3). Also the staining for the common GAMs marker Iba1 did not reveal any genotype-dependent differences in terms of glioma infiltration. However, we noted a striking genotype-dependent difference in the neutrophil influx into the glioma. Thus, infiltration of the MPO+ neutrophils was obvious in the gliomas in SorLA-KO brains, while it did not occur in the WTs (Fig. 4 E-F). This was not due to the overall increase in neutrophil amounts in the circulating blood of SorLA-KO mice, as the numbers of circulating neutrophils, as well as of erythrocytes, monocytes, lymphocytes, basophils, and eosinophils, were comparable in glioma-bearing WT and SorLA-KO mice (Fig. S5). We propose that this massive neutrophil infiltration is a direct consequence of a pro-inflammatory milieu promoting their migration into the brain parenchyma in SorLA-deficient mice. One of the key mechanisms facilitating neutrophils influx into the tissue involves TNFα-driven induction of ICAM1, an adhesion molecule critical for transendothelial migration of these cells (30, 31). Increased soluble ICAM1 (sICAM1) levels are also a well-established hallmark of inflammation (32). As anticipated from our data, sICAM1 was elevated in the tumor-containing hemispheres derived from SorLA-KO mice as comparted to the WTs (Fig. 4G). These results strongly supported our hypothesis that loss of SorLA shifts the properties of glioma microenvironment towards pro-inflammatory.

Finally, to further elucidate the mechanisms limiting tumor growth in SorLA-KO mice, we focused on cell death mechanisms that might be activated by TNFα itself, or by the infiltrating neutrophils. TNFα can trigger apoptosis or necroptosis via its receptor TNFR1 (33), while neutrophils elicit ferroptosis (34). We thus checked which of these pathway(s) are activated in glioma specifically in SorLA-KO mice. The levels of ferroptosis markers in glioma samples were not genotype-dependent. Moreover, we did not observe induction of apoptosis, as the cleavage of PARP and caspase-3 was negligible and similar for both WT and SorLA-deficient mice (Fig. S6). However, we noted increased levels of necroptosis markers p-RIPK3 and MLKL in the glioma-bearing hemispheres from SorLA-KO mice, as compared to the WTs (Fig. 4 H-I). These results suggested that necroptosis might contribute to the elimination of glioma cells and, in consequence, to limiting tumor growth in SorLA-KO brains.

## Discussion

SorLA is an important player in maintaining the functional integrity of the brain (3). Although this role has mainly been attributed to its neuronal functions in the past, single cell sequencing approaches discovered the complexity of SorLA expression patterns in the diseased brain. In particular, potential microglial activities of SorLA are gaining increasing attention due to relatively high expression levels of the receptor in this cell type (35, 36) and to the relevance of microglial activity to the pathogenesis of virtually all brain disorders (37). Here, we have focused on the microglial roles of SorLA in the context of glioma. However, it is plausible that such a function also bears relevance for other brain diseases, such as AD, and that the ability of SorLA to limit pro-inflammatory activity of microglia shown in this study may represent a mechanism of fundamental significance.

Here we discovered that the response of SorLA-KO mice to a developing glioma is shifted towards a pro-inflammatory state as compared to WT animals. This phenotype is manifested by morphological changes of microglial cells, increased sICAM1 levels, and by neutrophils infiltration. As a caveat, using a global SorLA-KO does not allow us to directly dissect functions of the receptor in GAMs versus other cells in the tumor microenvironment. In particular, while it is unlikely that altered neuronal activities in SorLA-KO mice directly shape the pro-inflammatory milieu, we cannot exclude that they contribute to limiting glioma growth via different mechanisms, as the activity of peritumoral neuronal cells may influence tumor growth and invasion (38, 39). Also lack of SorLA in neutrophils should be considered as a potential contributor to the observed phenotype. However, remarkable SorLA expression has not been reported *in vivo* in neutrophils (40–42), also not in the context of glioma (43). Importantly, lack of SorLA did not influence the numbers of circulating neutrophils in glioma-bearing mice, nor did it drive neutrophils infiltration to the tumor-free brain hemisphere. This observation strongly favors the hypothesis that neutrophils infiltration is a secondary consequence of the pro-inflammatory character of the tumor microenvironment in SorLA-KO mice.

We further propose that GAMs polarization towards the anti-tumor state in SorLA-KO brains is the plausible cause for the limited tumor growth. In our view, TNFα release from GAMs and neutrophils infiltration reflect the anti-tumor response that leads to such beneficial effects. Although the role of inflammation in tumor biology remains a matter of debate (44, 45), depleting TNFα in host cells resulted in larger tumors and shorter survival in the murine glioma model (46). In line with this notion, inhibition of Stat3 in a murine model of glioma enhanced TNFα expression in microglia/macrophages, blocked tumor growth and improved survival (47). Also in GBM patients, high level of the cytokine, both on the periphery and in the tumor microenvironment were linked to increased overall survival (48). Likewise, the role of neutrophils in tumor progression remains unclear, as their inflammatory activation has been linked to bad patient outcomes in several studies (49). However, recent study on the murine models of glioma suggests that neutrophils can limit glioma growth in the early stage of the disease (50). Neutrophils can undergo polarization into pro- or anti-tumor phenotypes (51), so it is tempting to speculate that the pro-inflammatory glioma microenvironment of SorLA-KO mice not only drives neutrophils recruitment, but more importantly, promotes their anti-tumorigenic functions.

In summary, unlocking the endogenous pro-inflammatory potential of microglia/macrophages directly at the site of glioma may represent a potent mechanism of limiting its growth. In this context, targeting *SORL1* expression in GAMs or the interaction between SorLA and TNFα emerges as an exciting strategy for the future pharmacological interventions in glioma. Further studies on the molecular details of the mechanism described herein will facilitate development of such novel therapeutic approaches.

## Materials and Methods

### scRNA-seq data analysis

The analysis covered 11 newly diagnosed glioblastoma (ndGBM) patients (6). The raw count matrices (GSE182109) were processed using the R package scTools (1.0). A list of samples was created with scCreateRawSamples(), from which cells expressing at least 3000 genes and genes expressed in at least 5 cells were selected. Standard quality control metrics were calculated with scIntroQC() and cells with mitochondrial read percentage < 15%, ribosomal read percentage > 5%, and red blood cell read percentage < 0.5% were kept. The filtered samples were integrated with the scIntegrate() function using the SCTransform algorithm (52). The most informative 3000 genes were selected to perform the Principal Component Analysis (PCA) and first 30 Principal Components were used for cell clustering with Uniform Manifold Approximation and Projection algorithm (UMAP) (53).

Markers of obtained UMAP clusters were found with scMarkers() that utilizes Seurat’s (54) FindAllMarkers() function, its criteria were: gene presence in at least 50% of cells, only positive fold-change (FC) and the logFC threshold equal to 0.25. Expert biological analysis was used to manually annotate glioma-associated microglia/macrophages cells (GAMs) that were used for further analysis.

Spearman correlation of *SORL1* expression with other genes using integrated counts from Seurat was calculated with the FDR correction for multiple testing (FDR<0.05). To select significant features and identify their non-linear relationships between discretized *SORL1* gene expression and other gene’s continuous expression, the MCFS-ID algorithm was applied (18, 19) on 75% of cells (training set). Remaining 25% cells were used to validate the predictive quality (using popular classifiers) of significant genes returned by MCFS-ID. Additionally, selected top genes from the same MCFS-ID returned ranking were visualized on the heatmap using pheatmap R package (55).

More details on scRNA-seq analysis are provided in the Supporting Information file. The source code related to the paper analysis is located in a GitHub repository: https://github.com/mdraminski/expressionLevelsSorLA.

### Cell lines

HEK293 and GL261 cells were cultured in DMEM (Gibco, #31885) with 10% fetal bovine serum (FBS, Gibco, #10500064), penicillin/streptomycin (Sigma, #P4333), with addition of 100 μg/mL G418 (Invivogen, #ant-gn-5) in case of GL261-tdTomato^+^Luc^+^ cells. BV2 cells were cultured in DMEM w/Glutamax (Gibco, #10569-010) with 2% FBS and penicillin/streptomycin. HEK293 transfections were performed with Lipofectamine 2000 reagent (ThermoFisher, #11668019) according to standard procedures and the cells expressing SorLA mutants and TNFα-GFP (or GFP) were collected 24 hours later.

### Primary microglia cultures

Primary microglial cultures were prepared using whole brains of P0-P1 C57BL/6J wild-type or SorLA-KO newborns as described (5) and cultured in DMEM w/Glutamax (Gibco, #10569-010) with 10% FBS (Pan Biotech, #P30-3302), and penicillin/streptomycin. In brief, glial cultures were plated on flasks coated with 0,1 mg/mL poly-L-lysine (Sigma, #P1274), washed 48 h after plating, and further cultured for 8 days prior to shaking-off microglial cells (1h, 80 RPM at 37°C). After centrifugation, microglial cells were seeded onto glass coverslips on 6- or 24-well plates (6 x 10^5^ or 1 ×10^2^ cells per well, respectively) and used for further procedures 48 h later.

### iPSC culture and differentiation to microglia cells

Human induced pluripotent stem cell (iPSC) line BIHi043A-Xmool and the *SORL1* deficient (SLKO) iPSC line were used. The SLKO iPSC line was generated by CRISPR/Cas9-mediated genome editing using the gRNA targeting exon 1 in the *SORL1* gene (5’ CAGTAGCGTTCGCCCGAACA ’3). SorLA deficiency in a clone containing a 8 bp frame shift deletion was confirmed by western blotting. iPSC cells were cultured on Matrigel (Corning, #356234)-coated plates in Essential 8™ Flex Medium (Gibco, #A2858501). Culture medium was changed daily and cells were passaged in clusters every 3-4 days at a density of 80% using 0.5 mM EDTA/PBS. iPSCs were differentiated to microglia as previously described (56). The detailed protocol is provided in Supporting Information. At day 38-42 the microglia were used for functional studies.

### Microglia stimulation

Primary microglia was stimulated with 100 nM phorbol myristate acetate (PMA, Sigma, #P8139) or 100 µg/ml lipopolysaccharide (LPS, Sigma, #L7770) in DMEM w/Glutamax (Gibco, #10569-010) for 24 h. For RNA isolation, cells were washed twice with cold PBS and collected. For cytokine levels analysis, cell medium was collected, centrifuged to remove cell debris, and frozen at −80°C until use. For immunostaining, glass coverslips with cells were fixed with 4% PFA in PBS. BV2 ells were plated on glass coverslips, stimulated with 100 nM PMA for 24 hours and fixed with 4% PFA in PBS.

For co-culture experiments, GL261 cells were seeded into the cell culture inserts with 0.4 µm pores (Falcon, #353090; 2.5×10^4^ cells/insert). Microglia cells were plated onto a 6-well plate (6×10^5^/well). After 24 h of medium conditioning over GL261 cells, medium and inserts were transferred to microglia-containing wells. After 24 h of co-culture microglial RNA was isolated.

iPSC-derived microglia were seeded at a density of 5×10^4^ cells per well in microglia differentiation medium in 96-well plates and stimulated with 1.0 µg/ml LPS (Sigma, #L4391) in microglia differentiation medium. After 24 hours, the medium was collected for further analyses and the cells were harvested for RNA isolation.

### ELISA measurements

Following enzyme-linked immunosorbent assay (ELISA) kits were used to measure cytokine levels in media samples: murine U-Plex Biomarker Group 1 assay (Meso Scale Diagnostics, #K15083K), human TNFa U-PLEX assay (MSD, #K151UCK). Mouse ICAM-1/CD54 Quantikine ELISA Kit (R&D Systems, #MIC100) was used to determine the levels of sICAM1 in tissue samples.

### Expression constructs

pEGFPC2-BIO plasmid used for GFP overexpression was a kind gift from Jacek Jaworski (57). GFP-TNFα was a gift from Jennifer Stow (Addgene plasmid # 28089; (58)). The SorLA-ΔCR construct (deletion of Cys1078 to Glu1552) has previously been described (59). The following constructs were generated: (1) SorLA-ΔVPS10P construct (deletion of Met1 to Pro753) in the pSecTag2A expression vector; (2) SorLA-ΔEGF/β-propeller construct (deletion of Leu754 to Glu1074) in the pcDNA3.1 expression vector; (3) SorLA-ΔFN3 construct (deletion of Val1555-Ala2123) in the pcDNA3.1 expression vector. Human SorLA mini plasmids were generated by amplifying the specific SorLA domain with overlapping primers using PCR technology. The fragments were then ligated into pcDNA3.1 zeo+ vector (Invitrogen). The *SORL1* stop codon was replaced with a myc-coding sequence. All mini receptors had the original SorLA signal and propeptide sequence and furin cleavage site at the N-terminus, and the original SorLA transmembrane and cytoplasmic domain at the C-terminus. The details on cloning strategy for deletion constructs are provided in the Supporting Information file. The information on SorLA domains covered by the constructs is provided in Table S5.

### Co-IP experiments

HEK293 cells overexpressing SorLA mutants and TNFα-GFP (or GFP) were lysed in 10 mM Tris pH 7.5, 150mM NaCl, 1mM CaCl2, 1mM MgCl2 and 0,5% Nondient P40 (NP40) substitute (Sigma, #11754599001) with proteases and phosphatases inhibitors. GFP-Trap Magnetic Particles® (Chromotek, #gtd) were used to pull down the GFP tag. Lysis buffer with reduced NP40 content (0.2%) was used for pull down and subsequent washing. Samples were boiled in 2x Laemmli Buffer to release bound proteins and examined by western blot.

### Quantitative RT–PCR

Total RNA was extracted from cell lysates and purified with RNeasy Mini Kit (Qiagen) or Direct-zolTM RNA MiniPrep Kit (Zymo Research). Reversely transcribed cDNA from total RNA was subjected to qRT–PCR using the following Taqman Gene Expression Assays: Sorl1 (Hs00983770; Mm01169526), β2M (Hs00187847; Mm00437762), Hprt1 (Mm00446968). Fold change in gene expression was calculated using the cycle threshold (CT) comparative method (2ddCT) normalizing to CT values for housekeeping gene (β2M for LPS and PMA stimulations, Hprt1 for co-culture with glioma cells).

### Human tissue samples

Human brain tissue samples used in this study were obtained from the archives of the department of Neuropathology of the Amsterdam UMC (University of Amsterdam, the Netherlands). Supporting Information Table S1 summarizes the clinical characteristics of patients. All human specimens were obtained and used in accordance with the Declaration of Helsinki and the Amsterdam UMC Research Code provided by the Medical Ethics Committee of the AMC.

### Mouse models

Mice with a targeted disruption of murine SorLA (SorLA-KO) have been described (60). We used SorLA-KO mice on an inbred C57BL/6J background. For primary cultures, SorLA-KO and C57BL/6J wild-type newborns of both sexes were used. For *in vivo* studies, SorLA-KO and C57BL/6J wild-type males at 10-16 weeks of age were used. Animals were kept under a 12 hr/12 hr light/dark cycle with free access to food and water. Animal experimentation was performed following approval by the First Ethical Committee in Warsaw (approvals no 1102/2020, 1274/2022).

### Stereotaxic implantation of glioma cells and bioluminescence imaging of tumor growth

GL261-tdTomato^+^Luc^+^ cells were stereotactically implanted into the right striatum of murine brains as previously described (61). Mice weight and wellbeing were controlled every 2-3 days. Luciferase activity of implanted GL261 tdTomato^+^Luc^+^ cells allowed for determination of tumor size using bioluminescence imaging. Measurements were performed at 7, 14 and 21 day post-surgery, as previously described (61). Detailed protocol is provided in Supporting Information.

### Separation of soluble fraction from brain lysates

21 days post glioma implantation brains of WT and SorLA KO mice were isolated, tumor-bearing hemispheres were dissected and homogenized in 20 mM Tris-HCl pH 7.5, 2 mM MgCl2 and 0.25 M sucrose, supplemented with protease and phosphatase inhibitors. Lysates were incubated on ice for 20 minutes and centrifuged (1000 g, 4°C, 10 min). Part of the resultant cleared lysate was kept for analyses, while the rest was further centrifuged (100 000 g, 4°C, 1 h) to obtain the soluble protein fraction. Protein concentration was measured using Pierce™ BCA Protein Assay Kit.

### Western blot

Tissue lysates were analyzed by western blot using the following antibodies: anti-myc (2278S, Cell Signaling), anti-SorLA (611861, BD Transduction Laboratories), anti-SorLA C-term (homemade, produced in rabbit), anti-GFP (SC8334, Santa Cruz), anti-pRIP3 (CS83613, Cell Signaling), anti-MLKL (ab66675, Abcam), anti-GAPDH (MAB374, Millipore), anti-caspase-3 (CS966, Cell Signaling), anti-PARP (CS9542, Cell Signaling), anti-TRFR (ab269513, Abcam), anti-GPX4 (ab125066, Abcam). Details are given in the Supporting Information (Extended Materials and Methods and Table S5). Signal was registered with the ChemiDoc Imaging System (Bio-Rad).

### Immunostaining

Following antibodies were used: anti-SorLA (MABN1793, EMD Millipore and homemade antibody produced in goat), anti-Iba1 (019-19741, WAKO), anti-TNFα (CS11948S, Cell Signaling), anti-vti1b (BD611404, BD Biosciences), anti-Rab11 (BD610657, BD Biosciences), anti-GM130 (BD610823, BD Biosciences), anti-Tmem119 (400002, Synaptic Systems), anti-MPO (AF3667, R&D Systems), anti-galectin3 (M3/38, Biolegend), anti-CD8alpha (ab217344, Abcam). Details on samples preparation and staining protocols are given in Supporting Information (Extended Materials and Methods and Table S5).

### Image quantification

Western blot signals were quantified using the Image Studio Lite software or Image Lab Software 6.1 (Bio-Rad). Microscopy images were quantified using Fiji software. Signal intensities were measured on single channel images as a mean gray value for the field of view. Colocalization analysis was performed with the Colocalization Threshold plugin with manual selection of the region of interest, containing the cell cytoplasm. Single z-plane images were used for calculating thresholded Mander’s coefficient (tM).

For the analysis of microglial cell morphology, 3D reconstruction of microglial branches revealed by Tmem119 staining was performed in Imaris 9.1.2. Software and Imaris Filament Tracer with spot detection mode. Reconstructed cell branches were subjected to Sholl analysis. More details are provided in the Supporting Information.

### Statistical analysis

For *in vivo* experiments, an indicated number n is the number of mice per group used in an experiment. For primary cell culture experiments, an indicated number n is the number of independent glial preparations (biological replicates) used for ELISA or qRT-PCR analysis. In case of colocalization studies, n is the number of individual cells quantified in a given experiment, and every experiment was replicated at least three times on independent microglial cultures. Statistical analyses were performed using GraphPad Prism software. Where applicable, outlier analysis was performed using Grubb’s test. The details of statistical analysis are specified in the figure legends.

## Acknowledgments

We are indebted to the Stem Cell Facility of the MDC for generation of the *SORL1*-deficient iPSC line and for providing microglia differentiation protocols. We thank Tatjana Pasternack and Beata Kaza for expert technical assistance. Part of the confocal imaging and 3D reconstructions were performed at the Polish Euro-BioImaging Node ‘Advanced Light Microscopy Node Poland’. This work was supported by the project financed by the Minister of Education and Science based on contract No 2022/WK/05, by the Foundation for Polish Science co-financed by the European Union under the European Regional Development Fund (Homing program, POIR.04.04.00 00 5CEF/18 00; to A.R.M.), by the National Science Center (OPUS program, 2020/37/B/NZ3/00761; to A.R.M.), and by the I.3.4 Action of the Excellence Initiative - Research University Programme at the University of Warsaw (to A.R.M.).

