## Supporting Information for "SorLA restricts TNFα release from microglia to shape glioma-supportive brain microenvironment"

*Anna R. Malik

**This PDF file includes:**

Supporting text: Extended Methods

Figures S1 to S6

Tables S1 to S5

Legends for Datasets S1 to S3

SI References

**Other supporting materials for this manuscript include the following:**

Datasets S1 to S3

### Supporting Information Text

### Extended Materials and Methods

#### scRNA-seq data analysis

Data acquisition and integration

Published dataset was used for the analysis (1). Only ndGBM (newly diagnosed glioblastoma) samples were selected, consisting of 11 patients (6 males and 5 females), some of whom had multiple samples taken (1). Downstream analysis of the raw count matrices obtained from the GEO series GSE182109 was performed using the R package scTools (1.0) (https://github.com/MateuszJakiel/scTools) with R version 4.2.0. A list of samples was created with scTools function scCreateRawSamples(), from which cells expressing at least 500 genes and genes expressed in at least 5 cells were selected. Then, using the scIntroQC() function the mitochondrial, ribosomal and red blood cell gene percentages were calculated. Only cells with mitochondrial read percentage < 15%, ribosomal read percentage > 5% and red blood cell read percentage < 0.5 were kept. Cells with gene counts that were less than 3000 were kept. These filtered samples were integrated using the scIntegrate() function. First, the SCTransform (2) algorithm was applied on all samples. The most informative 3000 genes were selected to perform the Principal Component Analysis (PCA) and first 30 Principal Components were used for cell clustering with Uniform Manifold Approximation and Projection algorithm (UMAP) (3).

Cell type marker calculation and cell type annotation

Markers of obtained UMAP clusters were found with scMarkers() that utilizes Seurat's (4) FindAllMarkers() function, its criteria were: gene presence in at least 50% of cells, only positive fold-change (FC) and the logFC threshold equal to 0.25. Expert biological analysis was used to manually annotate glioma-associated microglia/macrophages cells (GAMs) that were used for further analysis.

Data analysis

The integrated counts were extracted from the Seurat Object and were used to perform further calculations. The Spearman correlation of *SORL1* expression with other selected genes was calculated using cor.test(); p-values were adjusted using the FDR method with the FDR threshold 0.05. To detect non-linear relationships between *SORL1* expression and other genes, the MCFS-ID algorithm from the rmcfs R package (5, 6) was applied. *SORL1* gene expression was discretized by dividing the values of expression into three ranges: {(-3.1374856;  -0.3428585], (-0.3428585; 0.8406330], (0.8406330, +inf)} ie. ‘low’, ‘medium’ and ‘high’ levels. This discretized *SORL1* value was used as the response variable in the decision table and all other gene counts were used as explanatory features resulting in 38086 cells and 3000 genes. Next, 75% of cells from the input decision table were sampled to establish a training set, leaving the remaining cells for validation set. The significant features set returned by MCFS-ID was obtained using the permutation method and verified its predictive quality on the validation set using the following classifiers: decision tree, logistic regression, random forest, Naive Bayes and Support Vector Machines. Finally, to show differential levels of gene expression, in the context of discretized values of *SORL1*, selected top genes from the MCFS-ID returned ranking were visualized on the heatmap using pheatmap R package (7).

**iPSC culture and differentiation to microglia cells**

Human induced pluripotent stem cell (iPSC) lines were cultured on Matrigel (Corning, #356234)-coated culture plates in Essential 8™ Flex Medium (Gibco, #A2858501). Culture medium was changed daily and cells were passaged in clusters every 3-4 days at a density of 80% using 0.5 mM EDTA/PBS. iPSCs were differentiated to microglia using the described protocol (8). First, iPSCs were differentiated into hematopoietic progenitors (HPs) using STEMdiff™ Hematopoietic Kit (Stem Cell Technologies, #05310). On day -1, iPSCs with a density of 70-80% were passaged with ReLeSR™ (Stem Cell Technologies, #05872) into E8 flex containing Matrigel coated 6-well plates. Cell clusters of 100 cells were seeded at a density of 50-100 per well. At day 0, in wells with a total of 40-80 clusters, E8 flex medium was replaced with 2 ml medium A. On day 2, 1 ml medium A was added to the well. On day 3, the medium was removed and replaced with 2 ml Medium B. At day 5, 7, and 9 1 ml medium B was added to the well. At day 11, HPs were present in the media as well as attached to the bottom. To increase the yield of HPs in the collected media, the adherent cells were gently washed off using a 5 ml serological pipette. The cells were centrifuged at 300 x g for 5 min, resuspended in microglia differentiation medium (DMEM/F12, 2x Insulin-transferrin-selenite, 2x B27, 0.5x N2, 1x Glutamax, 1x non-essential amino acids, 400 µM monothioglycerol, 5 µg/ml Insulin, 100 ng/ml IL-34, 50 ng/ml TGFβ1, and 25 ng/ml M-CSF) and seeded at a density of $2x10^5$ cells in Matrigel (Corning, #354277)-coated 6 well plates (Corning). At day 13, 15, 17, 19, 21 1 ml of microglia differentiation medium was added to the well. At day 23, the medium, except 1 ml, was transferred to a 15 ml falcon tube to spin down the floating cells at 300 x g for 5 min. The cells were resuspended in 1 ml medium microglia diff medium and returned to the well. This was repeated for days 25-35. However, at day 35 the cells were resuspended in microglia differentiation medium plus 100 ng/ml CD200 and 100 ng/ml CX3CL1 to further mature the microglia. On day 37, 1 ml of the maturation medium was added to the well. At day 38-42 the microglia were ready for functional studies.

**Immunostaining procedures**

Mouse brains were dissected from tumor-bearing mice after transcardial perfusion with phosphate-buffered saline (PBS) and 4% paraformaldehyde (PFA) in PBS. After post-fixation (PFA, overnight) and cryopreservation in 30% sucrose/PBS, brains were cut in 40 µm free-floating coronal sections using a cryostat. Alternatively, the brains were dissected from non-perfused animals and slowly frozen on isopentane/dry ice. Next, 12 µm coronal sections were cut using a cryostat, immediately mounted on glass slides, and kept at –20°C until further use. Prior to staining, these sections were thawed at room temperature for 30 min and fixed in 4% PFA/PBS for 30 min. The sections were blocked in 1% horse serum in PBS and incubated with primary followed by secondary antibodies diluted in PBS, supplemented with 1% bovine serum albumin (BSA), 1% normal donkey serum (NDS) and 0.5% Triton-X. Specimens were washed with PBS, counterstained with DAPI and mounted with DAKO fluorescence mounting medium.

Cells grown on glass coverslips were fixed with 4% PFA/PBS. Next, the cells were washed with PBS, blocked for 1 hour in PBS supplemented with 5% NDS and 0.3% Triton-X, and incubated with primary antibodies followed by secondary antibodies diluted in PBS with 1% BSA and 0.3% Triton-X. After washing with PBS and counterstaining with DAPI, the coverslips were mounted with DAKO fluorescence mounting medium.

Human brain tissue samples used in this study for immunostainings were obtained from the archives of the department of Neuropathology of the Amsterdam UMC (University of Amsterdam, the Netherlands). Supporting Information Table S1 summarizes the clinical characteristics of patients. Human brain tissue was fixed in 10% buffered formalin, embedded in paraffin, sectioned at 5 μm, and mounted on pre-coated glass slides (Star Frost, Waldemar Knittel Glasbearbeitungs, Braunschweig, Germany).

Sections were deparaffinised in xylene, and ethanol (100%, 95%, 70%). Antigen retrieval was performed using a pressure cooker in 0.01 M sodium citrate buffer (pH 6.0) at 121°C for 10 min. Slides were washed with phosphate buffered saline (PBS; 0.1 M, pH 7.4) and incubated overnight with primary antibodies against SorLA (MABN1793, EMD Millipore, 1:150) and Iba1 (WAKO, 019-19741, 1:200) in antibody diluent (VWR International, Radnor, PA, USA) at 4 °C. The next day, sections were washed with PBS and incubated with Alexa Fluor 568 goat anti-rabbit or Alexa Fluor 488 donkey anti-mouse antibody (Invitrogen, Eugene, OR, USA, 1:200) plus Hoechst 33258 (1:1000; Thermo Fisher Scientific, Waltham, MA, USA) in antibody diluent (VWR International, Radnor, PA, USA) for 2h at room temperature, washed with PBS and mounted with Vectashield (Vector Laboratories Inc., Burlingame, CA, USA).

**Microglia morphology analysis**

For the analysis of microglial cell morphology, Z-stack images of WT and SorLA-KO murine brains after glioma implantation stained for Tmem119 were used. Each cell was selected manually using ImageJ software, and all stacks comprising it were duplicated and saved as a new file for further analyses. 3D reconstruction of microglial branches was performed in Imaris 9.1.2. Software. Imaris Filament Tracer with spot detection mode was used to determine starting and ending points. Automatically added ending points, which did not cover the cell surface were removed manually. Reconstructed cell branches were subjected to Sholl analysis.

**Murine model of glioma**

Stereotaxic implantation of glioma cells

Mice were kept under constant anesthesia during the whole procedure using 2% Isoflurane in oxygen. After administration of butorphanol (2 mg/kg body weight), meloxicam (2 mg/kg body weight) and bupivacaine (locally applied, 8 mg/kg body weight), skin on the head was incised. The hole in the skull was drilled at the following coordinates: -1 mm anterior-posterior (AP) and 2 mm medial-lateral (ML) from bregma. $80 \times 10^4$ of GL261 tdTomato+Luc+ cells in 1 µL of DMEM were stereotactically injected into the right striatum at the rate of 0.25 µL/min, to a depth of 3 mm according to the brain surface. Withdrawing of the syringe was performed at the rate 1 mm/min to avoid backward outflow of the cell suspension. The skin incision was closed using sutures. After surgery, animals were carefully monitored until they fully recovered from anesthesia. For the following 3 days, mice received meloxicam (2 mg/kg body weight) once a day. Mice weight and wellbeing were controlled every 2-3 days until the end of experiment.

Bioluminescence imaging of glioma growth

Mice implanted with GL261 tdTomato+Luc+ were injected intraperitoneally with D-Luciferin sodium salt (150 mg/kg body weight; Synchem, #BC218) in PBS. After 8 minutes, animals were anesthetized with 2% Isoflurane in oxygen and placed immediately in X-treme Imaging System (Bruker, Germany) under 2% isoflurane/oxygen supply. 10 minutes after D-Luciferin administration, bioluminescence emission was determined for the total time of 2 minutes. X-ray images were acquired following bioluminescence imaging. Tumors were visualized 7, 14 and 21 days post implantation. Bruker Molecular Imaging (MI) Software was used for signal quantification.

Blood sample collection

Prior to perfusions, samples of venous blood were collected immediately after incision of the right atrium. EDTA tubes (Profilab, #320) were used for material collection. Hematological analyses were performed by Vetlab® company, Warsaw, Poland.

A

B

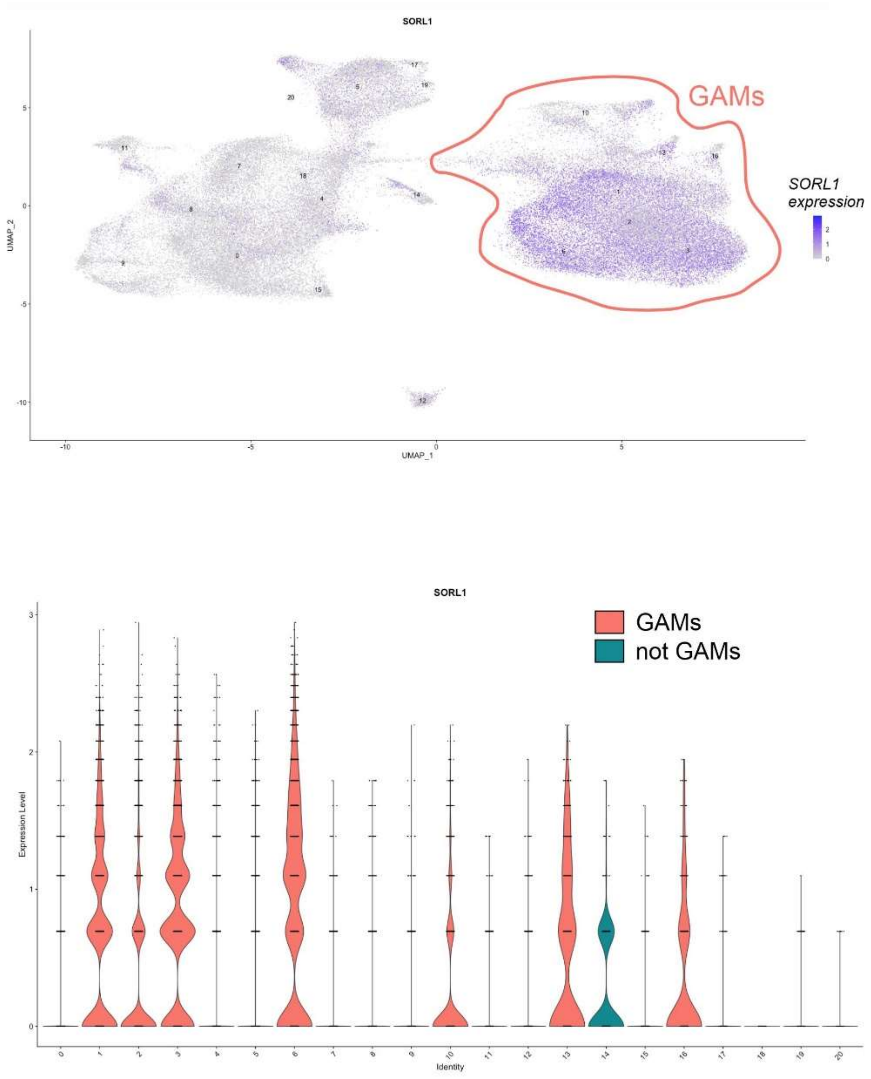


**Figure S1. Distribution of *SORL1* gene expression across single cells and analyzed clusters in ndGBM.**

(A) The UMAP plot displays the expression levels of *SORL1* in clusters of human GAMs, with SCT normalization used to represent the color intensity. The plot highlights clusters of cells containing GAMs in orange.

(B) The violin plot illustrates the expression of *SORL1* in all clusters present in the ndGBM data, where the orange color denotes clusters containing GAMs and green denotes those without GAMs. Clusters with the lowest *SORL1* expression (0,4,5,7,8,9,11,12,15,17-20) represent those without GAMs.

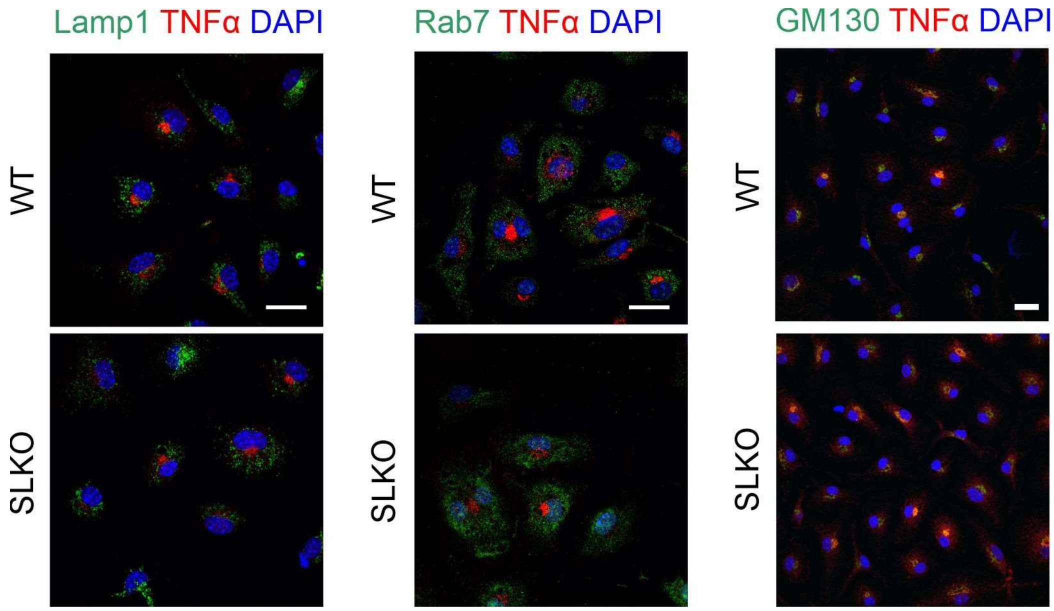


**Figure S2. Loss of SorLA does not influence colocalization of TNFα with lysosomes, late endosomes and Golgi in primary murine microglia.**

Representative images of primary murine WT and SorLA-KO microglia stimulated with PMA for 24h, immunostained for TNFα and the markers of lysosomes (Lamp1), late endosomes (Rab7) and Golgi (GM130). Cells were counterstained with DAPI. Scale bars, 20 μm.

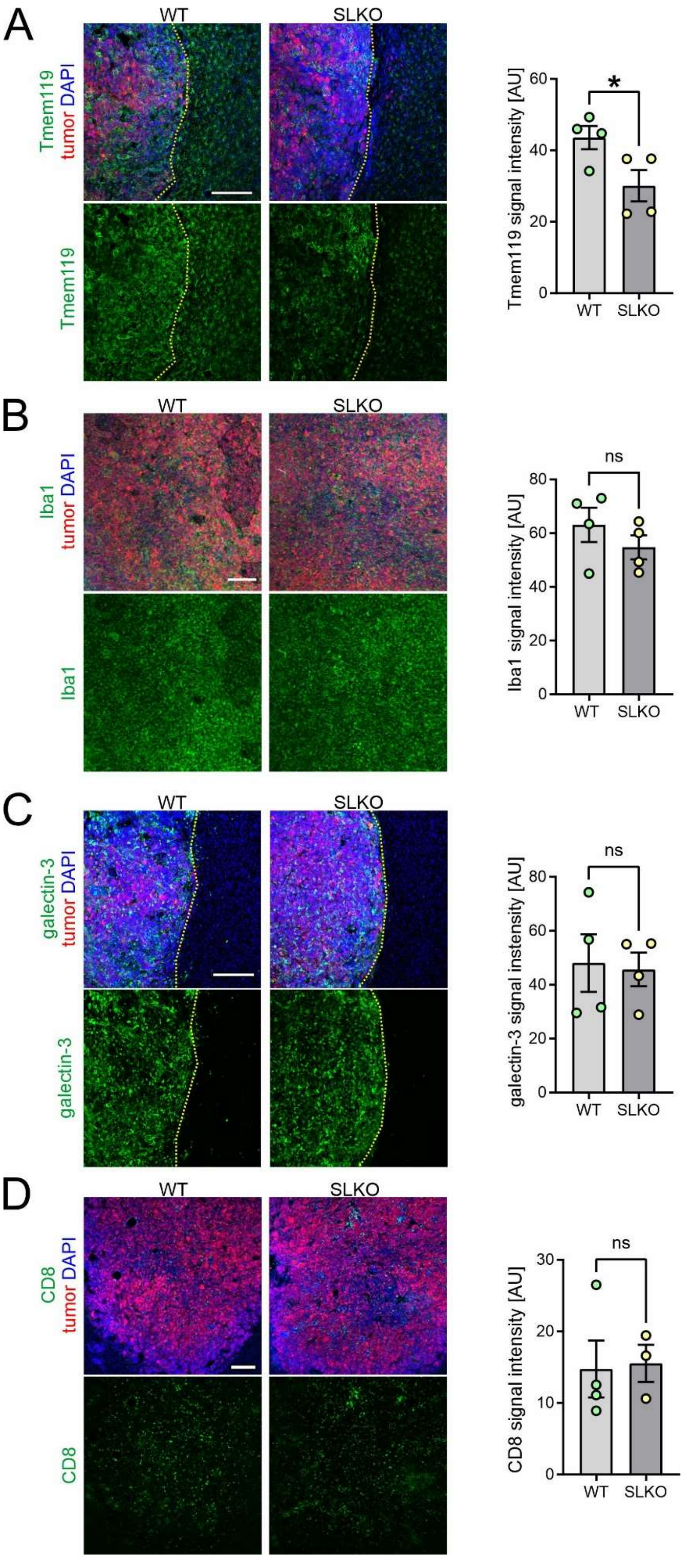
A
WT
SLKO
Tmem119 tumor DAPI
Tmem119
Tmem119 signal intensity [AU]
*
B
WT
SLKO
Iba1 tumor DAPI
Iba1
Iba1 signal intensity [AU]
ns
C
WT
SLKO
galectin-3 tumor DAPI
galectin-3
galectin-3 signal intensity [AU]
ns
D
WT
SLKO
CD8 tumor DAPI
CD8
CD8 signal intensity [AU]
ns

**Figure S3. Glioma infiltration by immune cells is unchanged in SorLA-KO mice.**

Representative images of the sections from glioma-bearing WT and SorLA-KO brains 21 days after implantation of GL261-tdTomato+Luc+ cells, immunostained for the markers of selected cell types. Tumor cells are seen in red. Sections were counterstained with DAPI (blue). Scale bars, 200 μm.

(A) Microglia marker Tmem119 is shown in green. Yellow dotted line marks tumor border. Mean signal intensity was quantified in the area outside the tumor. Mean ± SEM is shown. n=4 mice per genotype. *, $p<0.05$ in unpaired two-tailed t-test.

(B) Iba1, a marker of microglia/macrophages is shown in green. Mean signal intensity was quantified in the area inside the tumor. Mean ± SEM is shown. n=4 mice per genotype. ns, not significant in unpaired two-tailed t-test.

(C) Macrophages marker galectin-3 is shown in green. Yellow dotted line marks tumor border. Mean signal intensity was quantified in the area inside the tumor. Mean ± SEM is shown. n=4 mice per genotype. ns, not significant in unpaired two-tailed t-test.

(D) Cytotoxic T-lymphocytes marker CD8alpha is shown in green. Mean signal intensity was quantified in the area inside the tumor. Mean ± SEM is shown. n=3-4 mice per genotype. ns, not significant in unpaired two-tailed t-test.

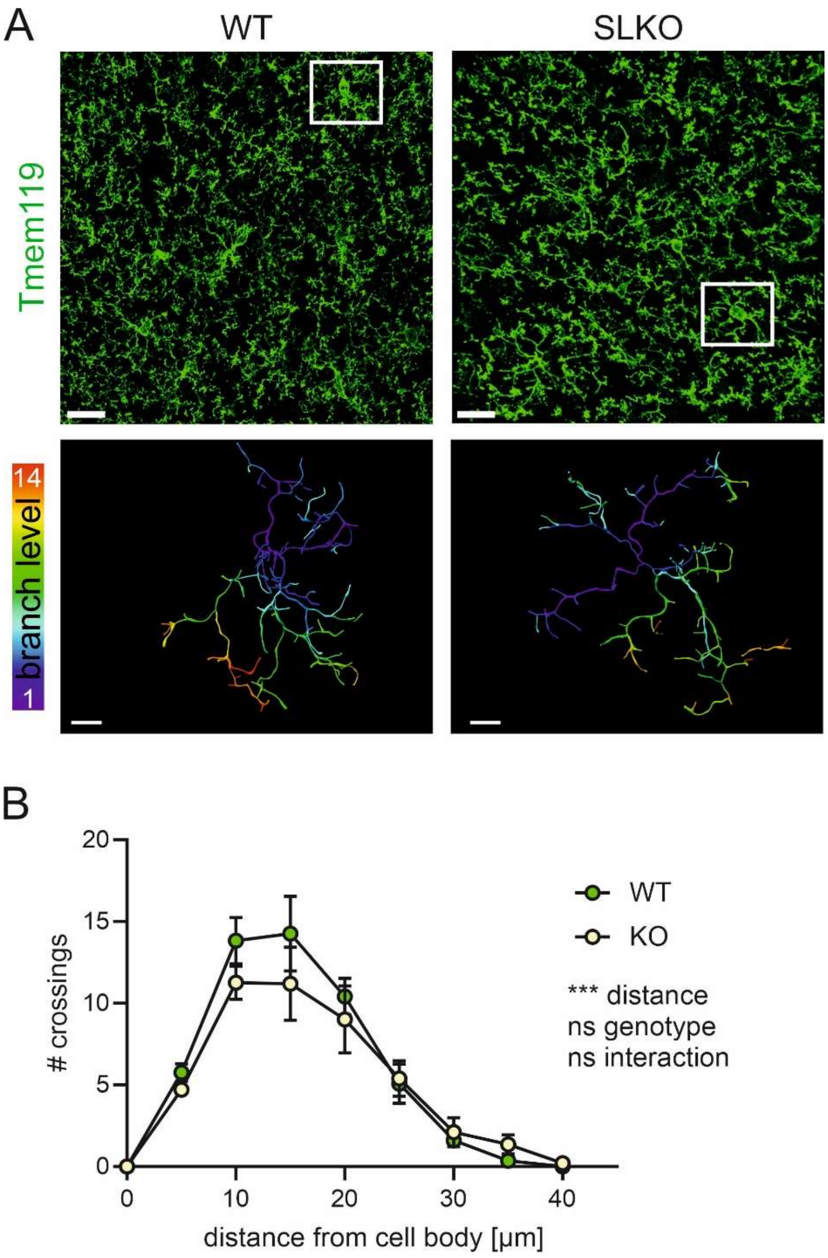


**Figure S4. Loss of SorLA does not alter microglia morphology in the contralateral hemispheres of tumor-bearing mice.**

(A) Upper panel: representative images of microglia morphology revealed by Tmem119 staining in WT and SorLA-KO mice in the contralateral hemispheres of glioma-bearing brains 21 days post-implantation. Scale bar, 20 µm. White box indicates the cell reconstructed below. Lower panel shows reconstructed microglia branches; color depicts branch level. Scale bar, 5µm.

(B) Sholl analysis of microglia morphology reconstructed as in (A). Mean ± SEM is indicated. ***, $p<0.001$; ns, not significant in two-way ANOVA; n=4 mice per genotype; for each mouse, 4-5 cells were quantified and an average of obtained values was treated as individual data point.

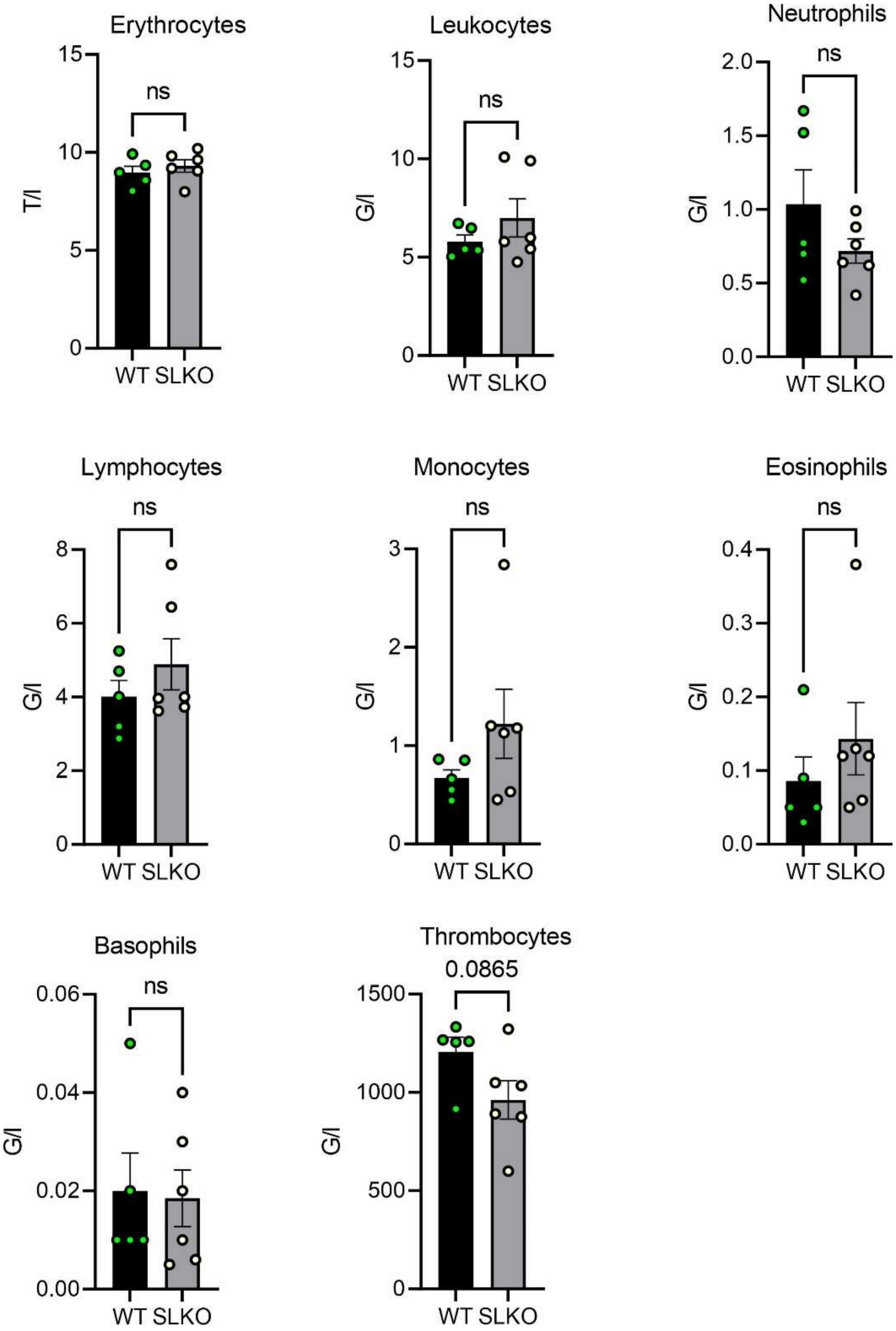


**Figure S5. Hematological analysis of WT and SorLA-KO glioma-bearing mice does not reveal significant differences in the numbers of blood cells between the genotypes.**

Erythrocytes, leukocytes, neutrophils, lymphocytes, monocytes, eosinophils, basophils and thrombocytes counts measured in the peripheral blood of WT and SorLA-KO glioma-bearing animals 21 days after implantation. Mean ± SEM is indicated. ns, not significant in unpaired t-test; n=5-6 mice per genotype.

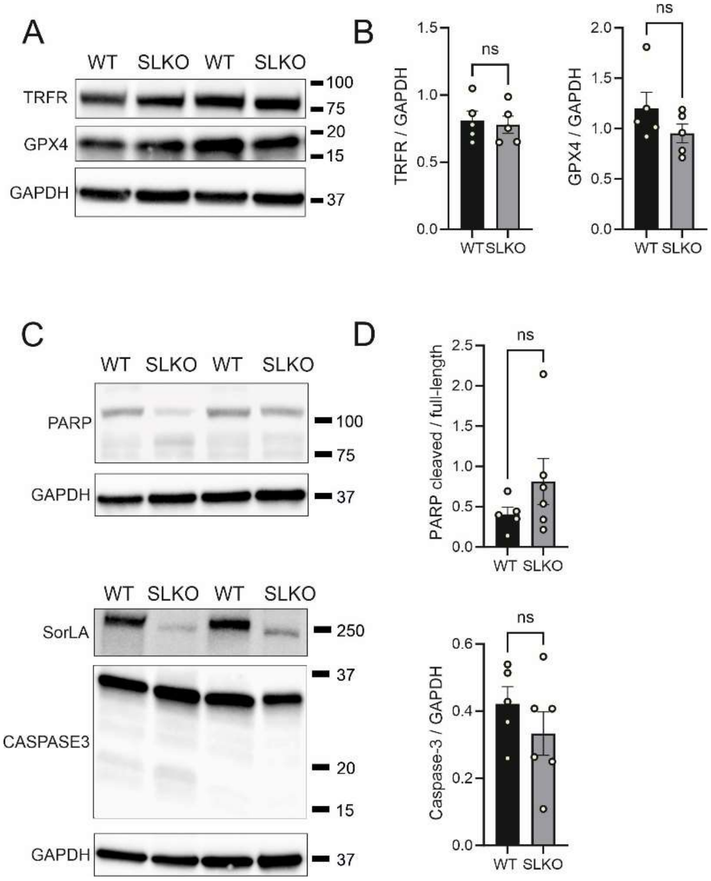


**Figure S6. Levels of ferroptosis and apoptosis markers are comparable in glioma-bearing hemispheres of WT and SorLA-KO mice.**

(A) Western blot analysis of ferroptosis markers, GPX4 and transferrin receptor (TRFR) in WT and SorLA-KO glioma-bearing hemispheres 21 days post-implantation. Detection of GAPDH was used as a loading control.

(B) Quantification of the western blot analysis as in (A). Signal intensities for GPX4 and TRFR were normalized to GAPDH signal. Mean ± SEM is indicated. ns, not significant in unpaired t-test; n=5 mice per genotype.

(C) Western blot analysis of apoptosis markers, caspase-3 and PARP in WT and SorLA-KO glioma-bearing hemispheres 21 days post-implantation. Detection of GAPDH was used as a loading control. Detection of SorLA is also shown. Caspase-3 and PARP are cleaved upon apoptosis activation which produces bands of lower size.

(D) Quantification of the western blot analysis as in (C). Ratios of signal intensities for cleaved and full-length PARP are shown. Signal intensity for full-length caspase-3 is normalized to GAPDH signal. Mean ± SEM is indicated. ns, not significant in unpaired t-test; n=5-6 mice per genotype.

**Table S1. Clinical characteristics of patients' samples and the summary of SorLA immunoreactivity in Iba1+ cells.**

| Patient # | Age, gender | tumor location | Pathology/diagnosis | SorLA expression in Iba1+ cells |
|---|---|---|---|---|
| 1 | 43, male | Parietal R | Astrocytoma IDH-wildtype, CNS WHO grade 3 | + |
| 2 | 63, male | Frontal R | Glioblastoma, IDH-wildtype, CNS WHO grade 4. | - |
| 3 | 11, female | Frontal L | Astrocytoma, IDH1-mutant, WHO CNS grade 4 | + |
| 4 | 56, male | Frontal R | Glioblastoma, IDH-wildtype, CNS WHO grade 4 | + |
| 5 | 17, male | Frontal R | Glioblastoma, IDH-wildtype, CNS WHO grade 4 | Hard to interpret |
| 6 | 16, male | Parietal R | Glioblastoma, IDH-wildtype, CNS WHO grade 4 | - |
| 7 | 64, male | Temporal R | Glioblastoma, IDH-wildtype, CNS WHO grade 4 | - |
| 8 | 50, male | Temporal R | Glioblastoma, IDH-wildtype, CNS WHO grade 4 | - |
| 9 | 49, female | Frontal R | Glioblastoma, IDH-wildtype, CNS WHO grade 4 | - |
| 10 | 54, female | Cerebellum | Glioblastoma, IDH-wildtype, CNS WHO grade 4 | + |
| 11 | 63, female | Parietal L | Glioblastoma, IDH-wildtype, CNS WHO grade 4 | + |
| 12 | 74, male | Fronto-parietal L | Glioblastoma, IDH-wildtype, CNS WHO grade | + |
| 13 | 48, male | Parieto-temporal L | Glioblastoma, IDH-wildtype, CNS WHO grade 4 | - |
| 14 | 11, male | Occipital L | Glioblastoma, IDH-wildtype, CNS WHO grade 4 | - |
| 15 | 40, male | Frontal L | Glioblastoma, IDH-wildtype, CNS WHO grade 4 | Hard to interpret |

**Table S2. Expression data extracted from Szulzewsky et al., Plos One, 2015** (9).

| glioma-associated vs naive CD11b+ cells | | | | | |
|---|---|---|---|---|---|
| transcript | AffyID | LogFC | FC | p.value | Adj.p.Val |
| SorCS2 | 10529515 | -1.0137488 | 0.4952577 | 2.13E-07 | 2.63E-05 |
| SorLA | 10592535 | 1.67434829 | 3.1917514 | 0.000794346 | 0.007875 |

**Table S3. Expression levels of GAMs marker genes in all cell clusters in newly-diagnosed GBM.**

| cluster | ***AIF1*** | ***CD68*** | ***ITGAM*** | ***P2RY12*** | ***TMEM119*** | ***CX3CR1*** | **GAMs score (Σ pct.1 values for all GAMs marker genes)** |
|---|---|---|---|---|---|---|---|
| 0 | n.a. | n.a. | n.a. | n.a. | n.a. | n.a. | n.a. |
| 1 | 0.929 | 0.800 | n.a. | n.a. | 0.588 | n.a. | 2.317 |
| 2 | 0.690 | 0.551 | n.a. | 0.725 | 0.540 | 0.59 | 3.096 |
| 3 | 0.821 | 0.708 | n.a. | n.a. | n.a. | n.a. | 1.529 |
| 4 | n.a. | n.a. | n.a. | n.a. | n.a. | n.a. | n.a. |
| 5 | n.a. | n.a. | n.a. | n.a. | n.a. | n.a. | n.a. |
| 6 | n.a. | n.a. | n.a. | 0.507 | 0.556 | 0.589 | 1.652 |
| 7 | n.a. | n.a. | n.a. | n.a. | n.a. | n.a. | n.a. |
| 8 | n.a. | n.a. | n.a. | n.a. | n.a. | n.a. | n.a. |
| 9 | n.a. | n.a. | n.a. | n.a. | n.a. | n.a. | n.a. |
| 10 | 0.731 | 0.674 | 0.569 | n.a. | n.a. | n.a. | 1.974 |
| 11 | n.a. | n.a. | n.a. | n.a. | n.a. | n.a. | n.a. |
| 12 | n.a. | n.a. | n.a. | n.a. | n.a. | n.a. | n.a. |
| 13 | 0.887 | 0.787 | 0.612 | n.a. | 0.796 | n.a. | 3.082 |
| 14 | n.a. | n.a. | n.a. | n.a. | n.a. | n.a. | n.a. |
| 15 | n.a. | n.a. | n.a. | n.a. | n.a. | n.a. | n.a. |
| 16 | 0.823 | n.a. | n.a. | n.a. | n.a. | n.a. | 0.823 |
| 17 | n.a. | n.a. | n.a. | n.a. | n.a. | n.a. | n.a. |
| 18 | n.a. | n.a. | n.a. | n.a. | n.a. | n.a. | n.a. |
| 19 | n.a. | n.a. | n.a. | n.a. | n.a. | n.a. | n.a. |
| 20 | n.a. | n.a. | n.a. | n.a. | n.a. | n.a. | n.a. |

pct.1 values for chosen marker genes in clusters 0-20 in ndGBM; data from Abdelfattah et al. Nat Commun, 2022 (1)

n.a. - given gene is not a marker gene for the particular cluster

Clusters 1, 2, 3, 6, 10, 13, 16 were classified as GAMs-containing clusters; these cells were pooled for further analysis of GAMs.

**Table S4. Antibodies used in the study.**

| Antigen | cat. number, manufacturer | purpose, dilution |
|---|---|---|
| SorLA | MABN1793, EMD Millipore | IF on human tissue, 1:150 |
| Iba1 | 019-19741, WAKO | IF on human and murine tissue, 1:200 |
| TNF | CS11948S, Cell Signaling | IF on cells, 1:200 |
| Sorla | homemade antibody, produced in goat | IF on cells, 1:100 |
| Vti1b | BD611404, BD Biosciences | IF on cells 1:100 |
| Rab11 | BD610657, BD Biosciences | IF on cells 1:200 |
| GM130 | BD610823, BD Biosciences | IF on cells, 1:1000 |
| myc-tag | 2278S, Cell Signaling | WB, 1:1000 |
| SorLA | Rabbit anti-SorLA (home-made; C-terminus) (10) | WB (Sorla deletion mutants) 1:1000 |
| SorLA | 611861, BD Transduction Laboratories | WB (full length SorLA), 1:1000 |
| GFP | SC8334, Santa Cruz Biotechnology | WB, 1:250 |
| Tmem119 | 400002, Synaptic Systems | IF on murine tissue, 1:500, |
| MPO | AF3667, R&D Systems | IF on murine tissue, 1:40 |
| galectin-3 | M3/38, Biolegend | IF on murine tissue, 1:500 |
| CD8-alpha | ab217344, Abcam | IF on murine tissue, 1:500 |
| p-RIP3 | CS83613, Cell Signaling | WB, 1:1000 |
| MLKL | ab66675, Abcam | WB, 1:250 |
| GAPDH | MAB374, Millipore | WB, 1:25 000 |
| caspase-3 | CS9662, Cell Signaling | WB, 1:1000 |
| PARP | CS9542, Cell Signalin | WB, 1:1000 |

| | | |
|---|---|---|
| TRFR | ab269513, Abcam | WB, 1:1000 |
| GPX4 | ab125066, Abcam | WB, 1:1000 |

WB, western blot; IF, immunofluorescence

**Table S5. SorLA domains covered by mini-receptors used in the study.**

| **SORLA domains** | **Amino acid number according to UniProtKB: Q92673.2** |
|---|---|
| Signal peptide | 1-28 |
| propeptide | 29-78 |
| Furin cleavage site | 79-82 |
| VPS10P | 124-755 |
| EGF/β-propeller | 756-1072 |
| CR | 1073-1550 |
| FN3 | 1551-2136 |
| transmembrane and cytoplasmic domain | 2138-2158 |

**Dataset S1 (separate file).** List of marker genes for cell clusters in ndGBM in a dataset from Abdelfattah et al., Nat Commun, 2022 (1)

**Dataset S2 (separate file).** List of marker genes for GAMs clusters.

**Dataset S3 (separate file).** Results of Monte Carlo Feature Selection analysis and further explanatory verification using Spearman correlation.
